# mir-21 is associated with inactive low molecular weight Argonaute complexes in thyroid cancer cell lines

**DOI:** 10.1101/2020.03.24.006072

**Authors:** Bonita H. Powell, Andrey Turchinovich, Yongchun Wang, Zhaohao Liao, Mohammad Aasif Dar, Gaspare La Rocca, George Essien Umanah, Martha A. Zeiger, Christopher B. Umbricht, Kenneth W. Witwer

## Abstract

Thyroid cancer is the most prevalent endocrine malignancy. We and others have shown that several microRNAs, which are post-transcriptional gene regulators, are aberrantly expressed in anaplastic thyroid cancer (ATC) and papillary thyroid cancer (PTC) tissues, as well as cell lines derived from these cancers. In the cell, miRNAs are bound to Argonaute (AGO) proteins as what could be termed low molecular weight RNA-Induced Silencing Complexes (LMW-RISCs) that can assemble with additional proteins, mRNA, and translation machinery into high molecular weight RISCs (HMW-RISCs) that exert regulatory function. In this study, we sought to analyze the association of miRNAs with RISC complexes in ATC and PTC. For ATC and PTC lines, miRNA species were enriched in both HMW-RISC and LMW-RISC cellular fractions, compared with intermediate molecular weight fractions and very low molecular weight (AGO-poor) fractions. Furthermore, 60% of all miRNAs were slightly more abundant in LMW-RISC versus HMW-RISC fractions by ~2-4 fold. Surprisingly, miR-21-5p, one of the most abundant miRNAs in both ATC and PTC lines and one of the most widely studied oncogenic miRNAs in many solid tumors, was consistently one the least abundant miRNAs in HMW-RISC and the most enriched miRNA in LMW-RISC fractions. These findings may suggest that miR-21 has a role or roles distinct from canonical post-transcriptional regulation in cancer. Furthermore, the methodology described here is a useful way to assess the distribution of miR-21 between HMW and LMW-RISCs and may help to reveal the true roles of this miRNA in thyroid cancer development, progression, and treatment.

## INTRODUCTION

Thyroid cancer is the most prevalent endocrine malignancy in the United States [1], [2]. Well-differentiated thyroid cancers (WDTC) such as papillary thyroid cancer (PTC) are the most common and typically have an excellent prognosis [3]. In contrast, undifferentiated tumors such as anaplastic thyroid cancer (ATC), are rare and almost invariably fatal [4], [5]. ATC can arise *de novo*, or more commonly, via dedifferentiation of WDTCs. Nevertheless, the precise mechanism whereby WDTC transitions to ATC is unknown [6] [7].

A host of aberrantly expressed microRNAs (miRNAs), e.g., miR-21, miR-146b, miR-221, and miR-222, have been extensively described in PTC and/or ATC [8] [9] [10][11][12][13][14]. Yet how these or other dysregulated miRNAs may influence thyroid tumor dedifferentiation remains incompletely understood. miRNAs are short, typically ~21-24-nucleotide (nt) non-coding RNA molecules that post-transcriptionally regulate gene expression by selectively interfering with messenger RNA (mRNA) translation [15][16]. In cells, mature miRNAs are complexed with Argonaute (AGO) proteins and may assemble with additional binding partners such as GW182 to form an RNA-induced silencing complex (RISC) [17][18][19][20][21]. An active RISC complex engages in post-transcriptional regulation when its miRNA recognizes a partially complementary sequence within a longer RNA: canonically, within the 3’ UTR of an mRNA [15], [22].

In cancers, including thyroid cancers, aberrant miRNA expression is commonly presumed to contribute to alterations in mRNA repression, which may promote pathogenesis [23] [24][25][26]. That is, miRNA and target RNA expression are presumed to be negatively correlated. However, this paradigm has been challenged in recent years [27]. Prominently, La Rocca et al. [28] reported that, in various adult mouse tissues *in vivo* and resting T-cells *in vitro*, and regardless of expression level, the vast majority of miRNAs are associated with AGO only. With few exceptions, just a small portion of AGO-bound miRNAs assembled into mRNA-associated RISCs [28]. These investigators also showed that RISC tended to be in a more active state in transformed cell lines, especially in the exponential growth phase and in mitogen-stimulated T-cells [28]. Because the activity state of RISC may be context-dependent, and miRNA activity could thus be uncoupled from overall abundance, we reasoned that a closer examination of active versus inactive miRNA species in thyroid cancer might provide valuable insights into pathogenesis.

In this study, we therefore separated cellular AGO-miRNA containing complexes using size exclusion chromatography (SEC) and profiled active and inactive AGO-miRNA in SW1736 (ATC) and TPC-1 (PTC) cell lines [28]. Active complexes, which comprise miRNA, AGO, GW 182, and mRNA, have a molecular weight that is normally > 2MDa [28]. Inactive complexes, with only AGO and miRNA, have a lower mass ~150kDa [28]. Hence, active and inactive complexes are referred to as high molecular weight RISC (HMW-RISC) and low molecular weight RISC (LMW-RISC), respectively (Figure. 1).

**Figure. 1.**
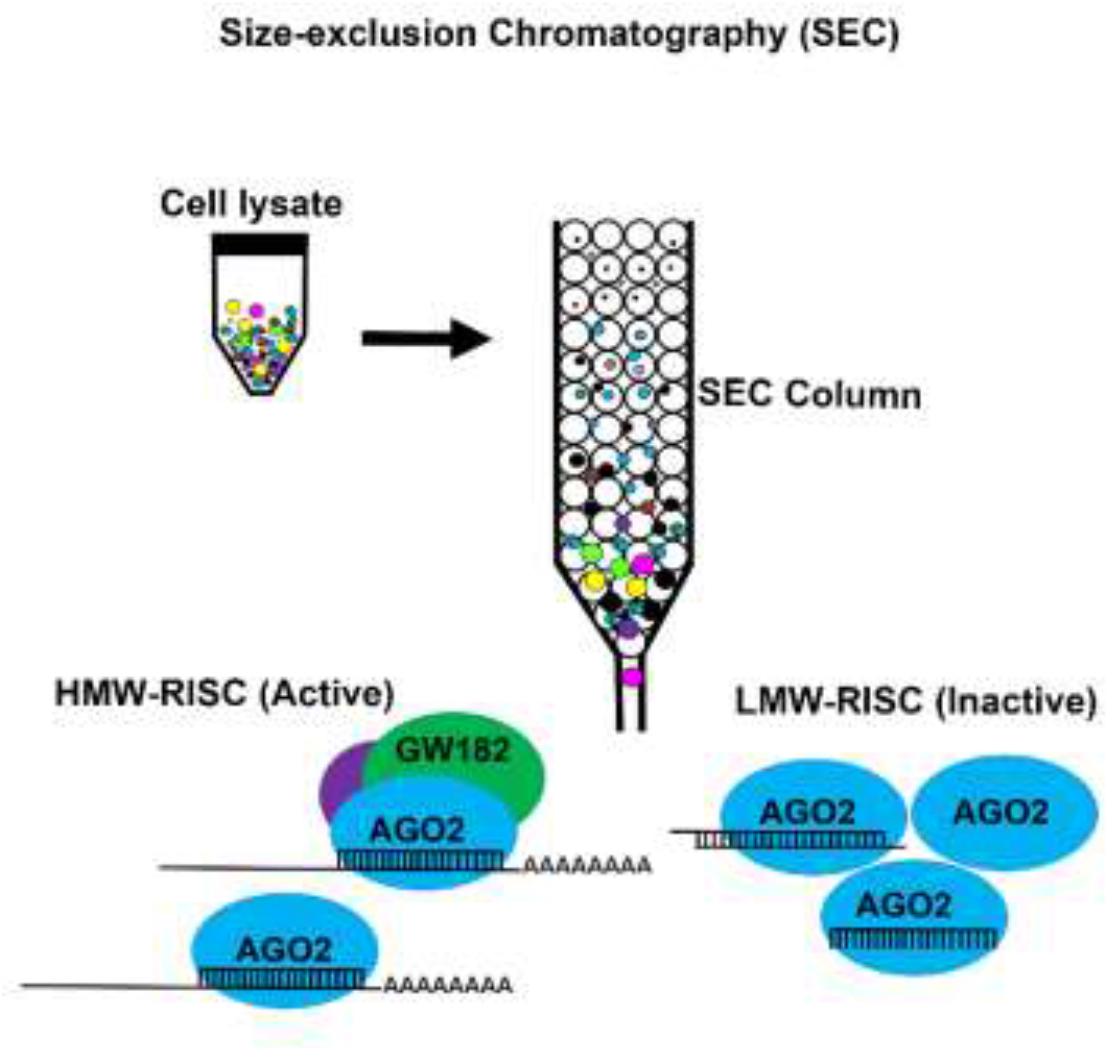
Illustration of RISC separation. Cell lysate is separated over a Superose 6 column. High molecular weight complexes including high molecular weight RISC (HMW-RISC) elute in early fractions, while low molecular weight fractions including Argonaute/miRNA complexes (LMW-RISC) elute later.

## MATERIALS & METHODS

### Cell Line Authentication and Culture

TPC-1 [29]and SW1736 [30] thyroid cell lines were authenticated by the Johns Hopkins Genetic Resources Core Facility by Short Tandem Repeat (STR) profiling. Cells were cultured in a humidified incubator at 37°C, 5% CO_2_ at 21% O_2_ (Normoxia) or 2% O_2_ (Hypoxia) using ProOx Compact O_2_/CO_2_ controller and chamber (Biospherix RS485). Cells were grown in RPMI −1640 medium (Gibco 11875-119) supplemented with: 10% heat-inactivated fetal bovine serum (FBS) (Sigma-Aldrich F4135-500 Lot#17E202), 2mM L-glutamine (Thermo Fisher 25030-081), 1% Non-Essential Amino Acids (Thermo Fisher 11140050), 10mM HEPES (Life Tech 5630-080), 100 U/mL Penicillin Streptomycin (Thermo Fisher 15140-122). Cells were seeded into five-layered tissue culture treated flasks (Corning cat#14-826-96) and grown until ~80% confluent.

### PBMC Isolation

Peripheral blood mononuclear cells (PBMCs) were isolated from fresh blood obtained from a human donor under a university IRB-approved protocol (JHU IRB #CR00011400). Up to 180mL of blood was collected into 3 60 mL syringes pre-loaded with 6 mL of anticoagulant acid citrate dextrose (ACD) (Sigma C3821 Lot#SLBW6172). Within 15 min of collection, whole blood was diluted 1:1 with 1X PBS containing 2% FBS, layered (appr. 35 mL per tube) onto 15 mL Ficoll-Paque PLUS (GE Healthcare 17-1440-02) using SepMate-50 tubes (Stem Cell Technologies 85450), and centrifuged for 15 min at 1200 x g. The plasma and buffy coat fractions were transferred to 50 mL conical tubes, diluted 1:1 with PBS, 2% FBS, and centrifuged for 8 min at 300 x g. The supernatant was decanted, and the cell pellet was suspended in red blood cell lysis buffer (155mM NH4Cl, 12mM KHCO3, 1mM EDTA, pH 7.3) and incubated for 10 min at 37°C. The suspension was diluted 1:1 with HBSS and centrifuged at 400 x g for 10 min. Cells were washed with PBS at 400 x g for 10 min. The cell pellet was snap frozen on dry ice and stored at −80°C.

### Transient miR-21-5p Mimic and Inhibitor Transfection

SW1736 cells grown in T-150 flasks (Corning #430825) to ~80% confluence were transfected with 10 nM of single-stranded miR-21-5p mimics (5’UAGCUUAUCAGACUGAUGUUGA-3’) (Integrated DNA Technologies IDT), 50 nM of duplex miR-21-5p mimics (Thermo Fisher Cat#4464066 ID#MC10206 Lot# AS02E016), 100 nM of miR-21-5p inhibitor (Thermo Fisher Cat#4464084 ID#MH10206 Lot#AS02DR3S) or negative control mimic (Thermo Fisher Cat#4464058 Lot#AS02CNS6) or inhibitor (Thermo Fisher Cat#4464076 Lot#AS02D9T1) using Lipofectamine 2000 (Invitrogen, Thermo Fisher cat#11668027) and Opti-MEM (Gibco, Thermo Fisher cat# 31985088). Block-iT Alexa Fluor control (Thermo Fisher 14750100) was used to assess delivery of RNAs. qPCR was done to optimize mimic transfection.

### Extract Preparation

~50×10^6 cells per line were harvested at 1000 x g for 5 min. Cell pellets were snap frozen on dry ice and stored at −80°C for 24 hrs. Cell pellets were resuspended in 600 μL freshly prepared ice-cold Sup6-150 buffer (150mM NaCl, 10mM Tris-HCl pH 7.5, 2.5mM MgCl2, 0.01% Triton X-100, 1X Halt phosphatase inhibitor Thermo Fisher 78428 Lot#TC260670, 1X Protease inhibitor Santa Cruz sc29131 Lot#D1618, 40U/mL Rnasin Promega N2115 Lot#0000310563, 1mM DTT Thermo Fisher20291 Lot#sk256023) and thawed on ice for 15 min. The lysate was vortexed for 30 s and cleared twice by centrifugation at 20,000 x g (5 min, 4°C). Lysate protein concentration was measured with BCA Protein Assay (Thermo Fisher 23252). Lysate was diluted to 4 mg/ml in Sup6-150, aliquoted, and stored at −80°C.

### Size Exclusion Chromatography

A Superose 6 column (GE Healthcare 17-5172-01 Lot# 10225796) was pre-equilibrated for 1.5 hrs at 4°C with Sup6-150 buffer without inhibitors or DTT on an AKTA Purifier system (GE Healthcare). 500 μL lysate (2 mg protein) was injected through a 0.75mm loop (GE Healthcare 18-1112-53). Run parameters were 0.5 mL/min and 1.5MPa for 1 column volume. Fractions 14-48 (0.5mL each) were collected; fractions 1-13 (column void) were discarded. 2 mg of gel filtration protein standard (Bio-Rad 151-1901) was run after sample collection to estimate approximate molecular weight of protein peaks. Half (250 μL) of each collected fraction was stored at −80°C for RNA extraction. The remainder was spiked with 500 ng of green fluorescent protein (GFP) (Thermo Fisher 13105S07E250) and concentrated to ~20 μL with 0.5 mL capacity 3kDa centrifugal filter units (Amicon/Millipore UFC500324) at 14,000 x g for 1.5 hrs at 4°C, then stored at −20°C in 1X Laemmli sample buffer (Bio-Rad 161-0747) with 10% betamercaptoethanol (BME) (Bio-Rad 161-0710) for < 1 week.

### AGO2-RNA Immunoprecipitation

Unfractionated input and pooled HMW-RISC, MMW-RISC, LMW-RISC, and RISC-Poor SEC fractions were incubated with 48μg of anti-AGO2 antibody (Sigma SAB4200085 Lot#087M4841V) for 16 hrs at 4°C with rotation. The mixture was then incubated with 18mg Protein G Dynabeads (Thermo Fisher 10003D) for 2 hrs at room temperature with rotation. Beads were washed 3X with ice-cold Sup6-150 and once with PBST, with beads retained by magnet at the end of each wash step. Bound protein was eluted with 240μL 50 mM glycine, pH 2.8. One volume of 1M Tris, pH 7.5 was added to the eluate. AGO2 was detected by Western blot and miR-21-5p by qPCR, using half of each sample for each assay.

### Western Blot

SEC fractions in loading buffer were heated to 95°C for 5 min and separated alongside a spectra multi-color protein ladder (Thermo Fisher 26634) on a 4-15% Tris-Glycine extended Stain-Free gel (Bio-Rad 5678085) at 100 V for 1.5 hrs using 1X Tris-Glycine SDS buffer (Bio-Rad 161-074). Gels were imaged with an EZ DocGel Stain-free imaging system (Bio-Rad 170-8274). Proteins were transferred to an absolute methanol-activated (10s) PVDF membrane (Bio-Rad 1620177) at 100 V for 1 hr in 4°C pre-chilled 1X Tris-Glycine buffer (Bio-Rad 161-0734). Membranes were blocked for 1 hr at room temperature in blocking buffer (PBS (Gibco 14190-144), 0.05% Tween20 (Sigma-Aldrich 274348500), 5% blotting-grade blocker (Bio-Rad170-6404)) and incubated with one or more of the following primary antibodies overnight at 4°C: AGO2 1:1500 dilution (Sigma-Aldrich SAB4200085 Lot#087M4841V), GW182 1:10,000 dilution (Bethyl A302-329A-T Lot#A302-329-T-2), PABP1 1:1000 dilution (Cell Signaling 4992 Lot#4), RPL26 1:1000 dilution (Bethyl A300-686A-M Lot#A300-686A-M-1), Actin 1:10,000 dilution (Sigma A1978-100μL Lot#065M4837V), GFP 1:1000 dilution (Roche 11814460001 Lot# 27575600). After three five-min washes with gentle rotation in blocking buffer, membranes were incubated with secondary antibody for 1 hr at room temperature: rat-HRP 1:10,000 dilution (abcam ab205720 Lot# GR32115155-1), rabbit-HRP 1:10,000 dilution (Santacruz sc-2004 Lot#B2216), or mouse-HRP 1:10,000 dilution (Santacruz sc-516102 Lot#c1419). Membranes were washed 3X in blocking buffer, then 2X in PBS with 0.05% Tween20, and incubated with Super Signal Chemiluminescent Substrate (Thermo 34580) for 5 min with gentle rotation and imaged by iBright FL1000 (Invitrogen).

### RNA Extraction and qPCR

Total RNA was extracted from each SEC fraction by miRNeasy serum/plasma kit (Qiagen 1071073 Lot#160020206) after adding 1.0 x 10^8 copies/μL of cel-miR-39 exogenous spike-in control (Qiagen 219610 Lot#157036035), according to the manufacturer’s instructions. RNA concentration and purity were measured by NanoDrop (Thermo Fisher) spectrophotometer. Total RNA and small RNA quality control was by Fragment Analyzer (Agilent version# 1.1.0.11). qPCR was performed with 2 μL RNA input from each fraction, or from pooled fractions HMW-RISC (15-16), MMW-RISC (17-26), LMW-RISC (27-35), and RISC-Poor (36-47) using TaqMan microRNA Reverse transcription kit (ABI 4366597 Lot#00636931), TaqMan miRNA stem-loop RT primers/qPCR primer probe sets (ABI 4427975) (cel-miR-39 ID# 000200 Lot#P180110-003B10, miR-16 ID# 000391 Lot#P171018-000H05, U6 ID# 001937 Lot#P170729-001405), TaqMan Universal Master Mix II, no UNG (ABI 4440040 Lot#1802074), as described in manufacturer’s protocol, on a CFX96 Real-time System (Bio-Rad). For each RNA fraction, miRNAs and U6 were normalized to cel-miR-39, then to fraction 14 using the 2^-ΔΔCT^ method followed by log2 transformation Pooled RNA fractions (HMW-RISC, MMW-RISC, LMMW-RISC, and RISC-Poor) were normalized to unfractionated/input then to RISC-Poor by the 2^-ΔΔCT^ method followed by log2 transformation [31].

### Small Library Preparation and Sequencing

Unfractionated input, HMW-RISC, MMW-RISC, LMW-RISC, and RISC-Poor small sequencing libraries were constructed with CATS RNA-seq kit (Diagenode C05010041 Lot#4). Library primer dimers were removed by AMPureXP beads (Beckman). Quality control was assessed by Bioanalyzer high sensitivity assay (Agilent). Libraries of 160-180 bp were size-selected by bluepippin (Sage Science). Libraries were spiked with 20% PhiX Control v3 (illumina 1501766) and sequenced using Next Seq with 1 × 50 Mid output of 130M reads by the Johns Hopkins Microarray and Deep Sequencing Core. Sequencing quality control was performed by FASTQC (Babraham Bioinformatics) followed by adapter and polyA trimming with Cutadapt 1.17. Reads were sequentially aligned to reference transcriptomes rRNA, tRNA, RN7S, snRNA, snoRNA, scaRNA, VT-RNA and Y-RNA. All reads that did not map to the aforementioned RNAs were aligned to pre-miRNA, protein-coding mRNA, and long non-coding RNAs (lncRNAs). Species with low read counts were filtered out such that average number of reads for a gene in all samples were >=5. For each fraction, all reads that did not map to the human transcriptome were aligned to the hg38 genome reference (rest_hg38). Hg38_reads were used to calculate log2 [counts-per-million+1] for each gene. The log2[cpm+] difference between different conditions was used as a final metric to determine relative abundance. Sequencing data were deposited with the Gene Expression Omnibus (GEO; database accession number GSE146015). See supplementary section for bash scripts.

## RESULTS

### RISC and Associated Proteins Across Molecular Weight Fractions

SEC was performed with protein lysates from cell lines SW1736 (ATC) and TPC-1 (PTC) to separate active and inactive RISCs by molecular weight (Figure. 1). Enrichment of AGO2, a key component of RISC, as assessed by Western blot (WB), was used to define the SEC elution profiles of active and inactive complexes [28]. For both cell lines, AGO2 was detected in elution fractions 15-37, but was highly enriched in high molecular weight (HMW) fractions 15&16 (>2MDa) and low molecular weight (LMW) fractions 27-35 (~150kDa-100kDa) (Figure. 2A). Additionally, presence of fully-assembled, presumably functional RISC was further determined by enrichment of GW182, PABP, and RPL26. As previously demonstrated by La Rocca et al., each RISC-associated protein was present chiefly in HMW fractions 15-16 (Figure. 2A) (Supplementary Figure. 1) [28]. Because active RISCs predominantly eluted in HMW fractions 15-16, and inactive AGO-miRNA complexes eluted in LMW fractions 27-35, fractions 15 and 16 are referred to as high molecular weight RISC (HMW-RISC) and 27-35 as low molecular weight RISC (LMW-RISC) (Figure. 2A).

**Figure. 2.**
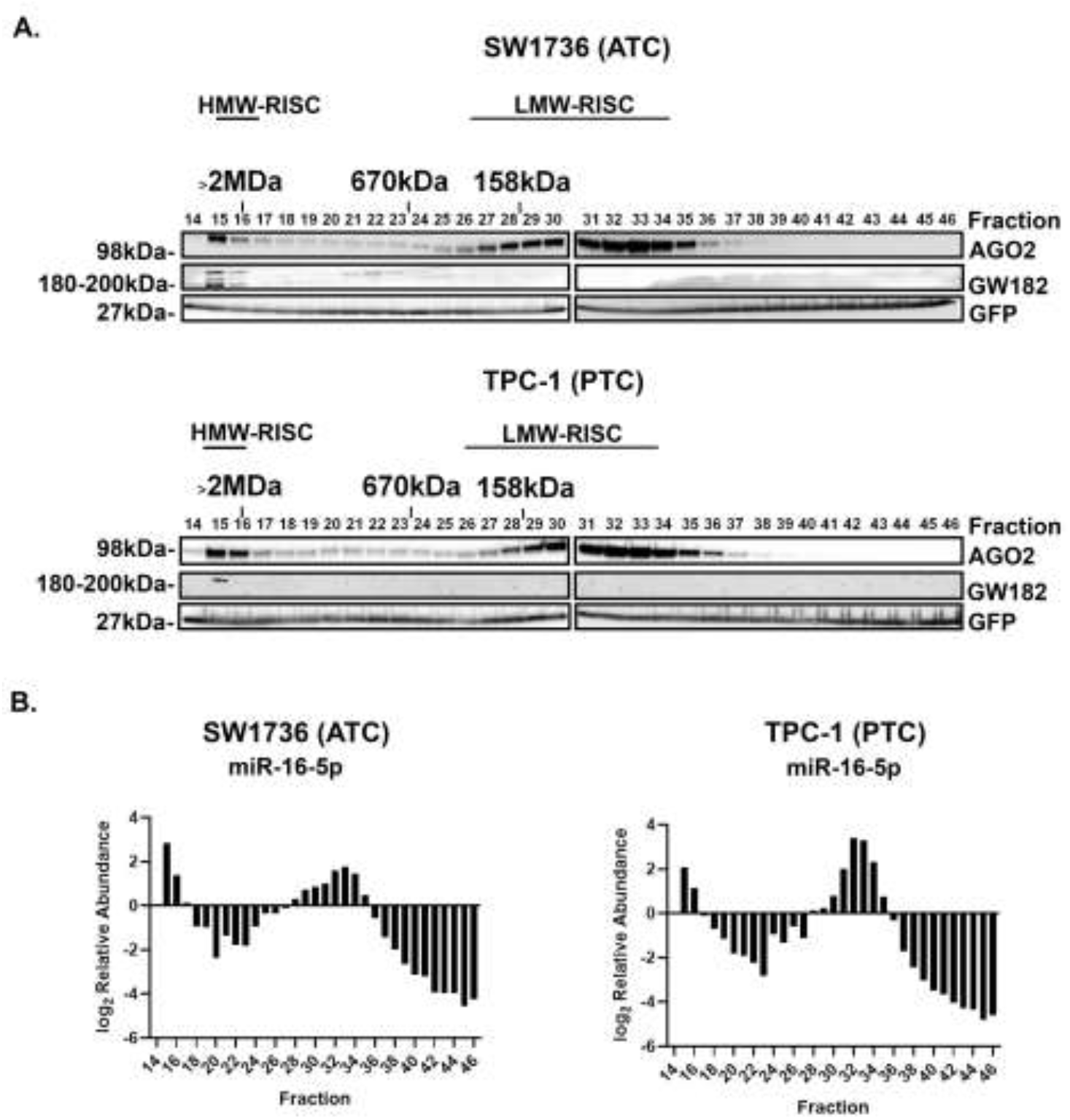
Protein and miRNA distribution across SEC fractions. **A.** SW1736 and TPC-1 mapping of protein content (Western blotting) across size exclusion fractions 14-46: AGO2 (~98 kDa) and GW182 (18Q-2QQ kDa), plus a green fluorescent protein (GFP) spike-in control (27 kDa). **B**. SW1736 and TPC-1 miR-16 content across SEC fractions 14-46, as assessed by qPCR.

### Representative Small RNA Distribution Across Molecular Weight Fractions

miR-16, an abundant miRNA in most cell types and often used as a normalization control, was evaluated in each SW1736 and TPC-1 SEC fraction by RT-qPCR. miR-16 was detected in all fractions but was present predominantly in high molecular weight fractions 15 and 16 (>2MDa) and low molecular weight fractions 27-35 (~150kDa), respectively (Figure. 2B). This distribution closely resembled the elution profile of AGO2. In contrast, U6 small nuclear RNA (U6 snRNA), which is not known to complex specifically with RISC proteins, was enriched chiefly in SEC fractions 20-30, rather than in the AGO/miRNA-rich fractions (Supplementary Figure. 2). This result suggests that miR-16 elution across molecular weight fractions mirrors that of AGO and may be AGO-dependent.

### HMW-RISC Fractions are rRNA- and mRNA-Enriched

miRNAs associated with HMW-RISC and LMW-RISC fractions in SW1736 and TPC-1 cells were next profiled more broadly by sequencing pooled HMW and LMW AGO2/miR-16-rich fractions. Additionally, fractions 17-26, which are presumably “intermediate” or “medium” RISCs (MMW-RISC), and fractions 36-47, which have very low levels of AGO2 and miR-16 (RISC-poor), were also pooled and sequenced. Ligation-independent small RNA sequencing, which enables analysis of RNA fragments of all classes regardless of end modifications, was performed using each of the pools. Relative to other RNA classes, pre-miRNA and mature miRNA comprised <1% of total RNA in each fraction, in agreement with a previous report [32]. Ribosomal RNA (rRNA), which was well represented in each fraction, had the highest mapping percentage in HMW-RISC and MMW-RISC fractions (Figure. 3). This is consistent with the association of active RISC with polysomes. We corroborated this by analyzing ribosome marker (RPL26) enrichment in HMW-RISC and MMW-RISC fractions (Supplementary Figure. 1) [28]. HMW-RISC and MMW-RISC fractions, as expected, also showed significant mapping to protein coding mRNA (Figure. 3). The mRNA transcripts present in LMW-RISC and RISC-poor fractions were also shorter than fragments detected in HMW-RISC and MMW-RISC. For instance, mRNA transcripts such as CTC1, TTC39B and PDPR had a much lower mapping coverage in LMW-RISC fractions compared to transcripts associated with the heavier RISCs (Supplementary Figure. 3–5). In contrast, tRNA fragments (tRNAFs), which may be associated with AGO, were highly enriched in LMW-RISC and RISC-poor fractions (Figure. 3) (Supplementary Figure. 6) [33]. Taken together, the HMW-RISC fractions associate preferentially with rRNA and mRNA compared with LMW-RISC fractions.

**Figure. 3.**
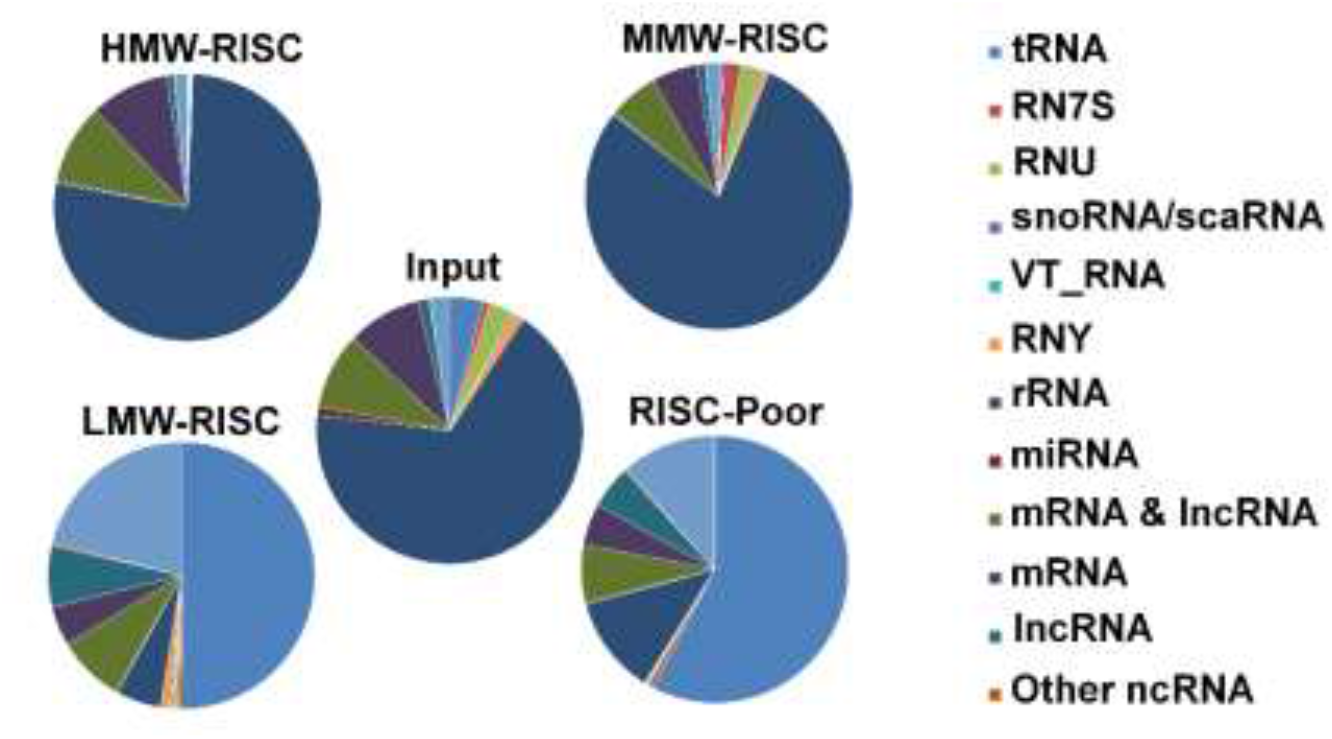
RNA class mapping percentages: RNA sequencing. Small RNA sequencing was performed for SW1736 and TPC-1 RNA from unfractionated input and pooled HMW-RISC, MMW-RISC, LMW-RISC, and RISC-poor SEC fractions. Depicted are the percentages of reads mapped to indicated classes of RNA.

### SW1736 and TPC-1 miRNAs Exist Predominantly in Low Molecular Weight Fractions

As seen with miR-16 enrichment across the individual SEC fractions (Figure. 2B), SW1736 and TPC-1 miRNAs detected by sequencing were most abundant in the pooled HMW-RISC and LMW-RISC fractions when compared with MMW-RISC and RISC-poor fractions (Supplementary figure. 7). Of 101 miRNAs that were abundantly expressed in both SW1736 and TPC-1 cells, approximately 60% of total miRNA species had greater enrichment in LMW-RISC fractions relative to HMW-RISC (Figure. 4 A). This included miRNA miR-21-5p, which was the most abundant miRNA in LMW-RISC fractions in each cell line [34] (Figure. 4 B).

**Figure. 4.**
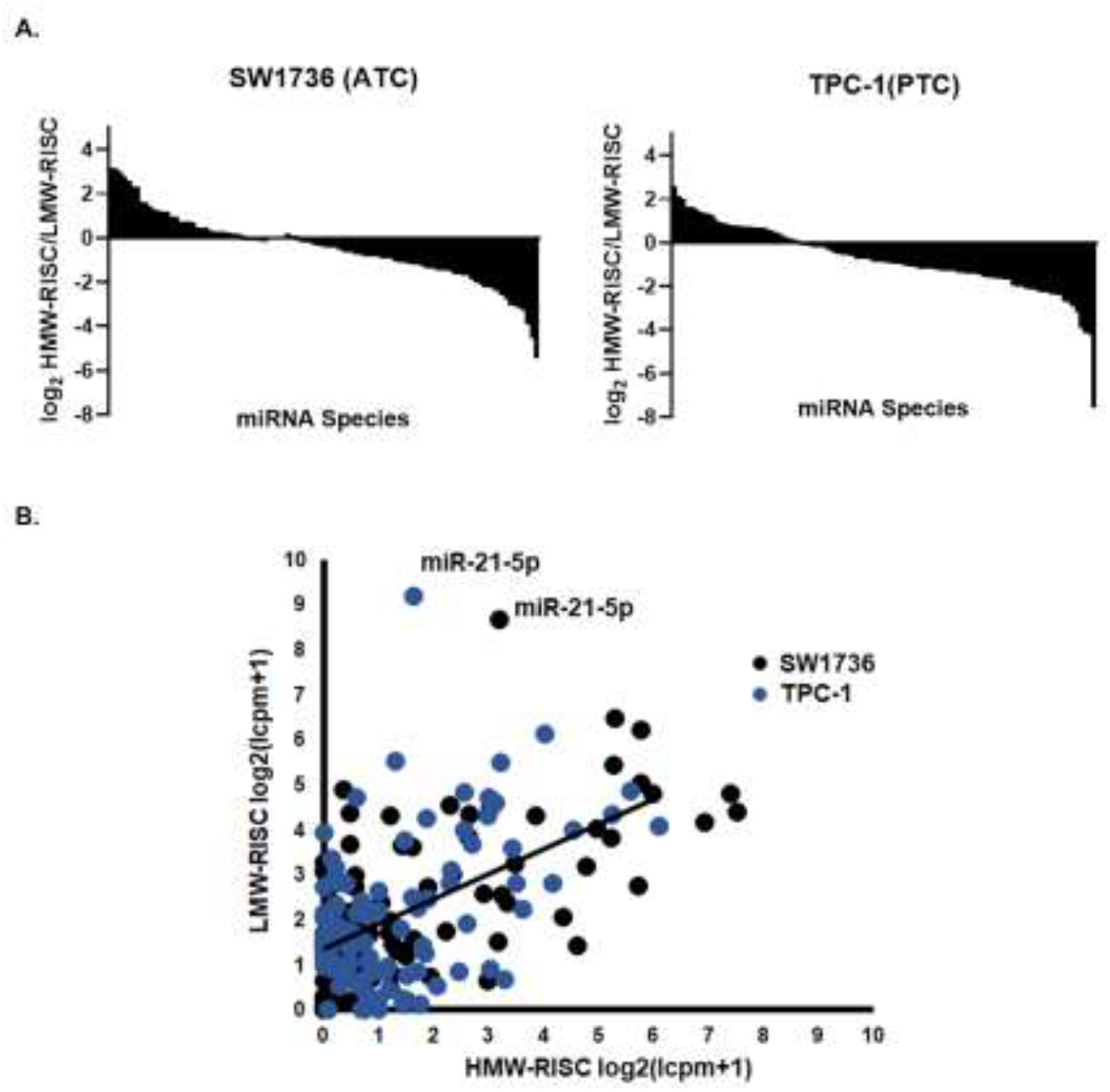
miRNA abundance in HMW-RISC versus LMW-RISC for all miRNAs detected in both fractions. **A**. HMW-RISC: LMW-RISC ratio of 101 miRNAs for both SW1736 (ATC) and TPC-1(PTC). **B**. Plot of (log2 counts per million + 1, lcpm+1) in HMW-RISC versus LMW-RISC for individual miRNAs shown in A, emphasizing miR-21-5p as an outlier enriched in LMW-RISC of both SW1736 and TPC-1.

### miR-21-5p is Differentially Enriched Between SW1736 and TPC-1 in LMW-RISC Fractions

Despite having a similar miRNA distribution across SEC fractions, the magnitude of individual miRNA enrichment in HMW-RISC or LMW-RISC fractions varied between SW1736 and TPC-1. For instance, miR-16 HMW-RISC enrichment was greater in SW1736 than in TPC-1 by 3-fold, while LMW-RISC enrichment was greater in TPC-1 by 7-fold (Figure. 1B). Likewise, miR-21-5p, which was the most highly abundant miRNA present in SW1736 and TPC-1 LMW-RISC fractions, was enriched in TPC-1 LMW-RISC fractions more than 2-fold greater than in SW1736 fractions (Figure. 4 B).

The HMW-RISC and LMW-RISC distribution of miR-21 compared with other dysregulated ATC and PTC miRNAs, including miR-146b, miR-221 and miR-222, was also measured by qPCR analysis of pooled RISC fractions (Figure. 5A). Of the four miRNAs, differential enrichment between SW1736 and TPC-1 was observed only for miR-21-5p, which was more enriched in LMW-RISC fractions by > 2-fold (Figure. 5A).

**Figure. 5.**
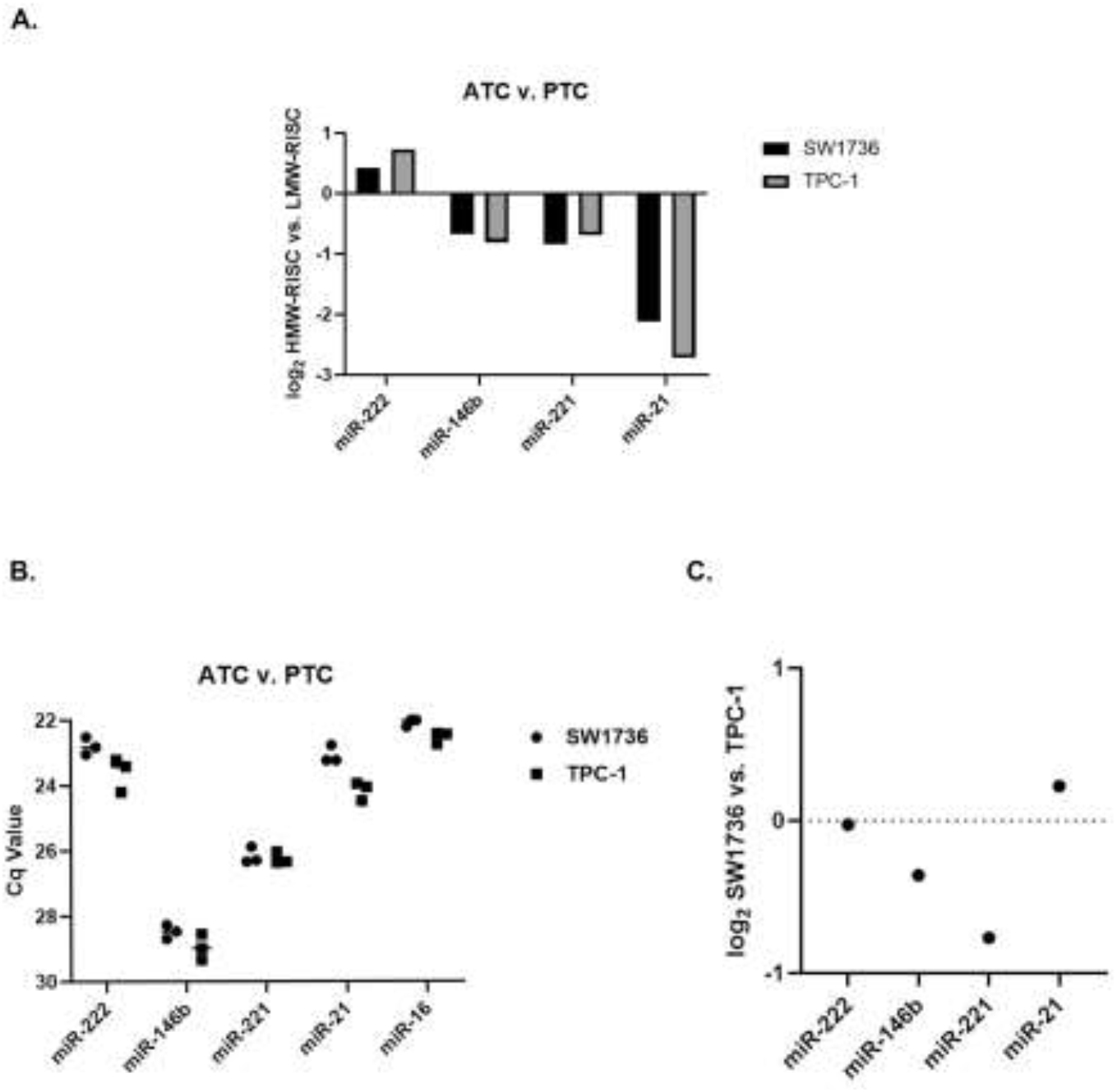
Individual qPCR assay verification of miRNA distribution and abundance. **A**. qPCR analysis of selected miRNAs miR-222-3p, miR-146b-5p, miR-221-3p, and miR-21-5p in pooled HMW-RISC and LMW-RISC fractions normalized to RISC-poor fractions in SW1736 and TPC-1. **B**. Abundance as indicated roughly by raw Cq values of miR-222-3p, miR-146b-5p, miR-221-3p, and miR-21-5p. **C**. Abundance ratio (data normalized to miR-16-5p) of dysregulated ATC and PTC miRNAs (SW1736 versus TPC-1).

Additionally, the relative expression levels of these miRNAs between SW1736 and TPC-1 were compared by qPCR. Confirming the results of RNA-seq, qPCR showed that the profound enrichment of miR-21-5p in LMW-RISC cannot be explained simply by overall abundance of miR-21-5p compared with other miRNAs, as miR-21-5p is not the most abundant miRNA in these cell lines. Furthermore, when each miRNA was normalized to miR-16 or U6-snRNA, all differences between SW1736 and TPC-1 were < 2-fold (Figure. 5C). This implies that the relative distribution of a given miRNA may be cell type-specific and independent of expression levels.

### AGO2-RIP Confirms AGO2-Bound miR-21-5p Enrichment in LMW-RISC Fraction

Although the elution profiles of AGO2 and miRNAs were similar, AGO2-RNA immunoprecipitation (AGO2-RIP) of pooled RISC SEC fractions was performed to confirm that miRNAs enriched in HMW-RISC and LMW-RISC fractions were bound to AGO2. Like the individual SEC fractions, AGO2 was most abundant in TPC-1 HMW-RISC and LMW-RISC fractions after AGO2-RIP and WB (Figure. 6A). Further, AGO2-RIP followed by RT-qPCR confirmed that LMW-RISC enriched miR-21-5p is complexed with AGO2 (Figure. 6B).

**Figure. 6.**
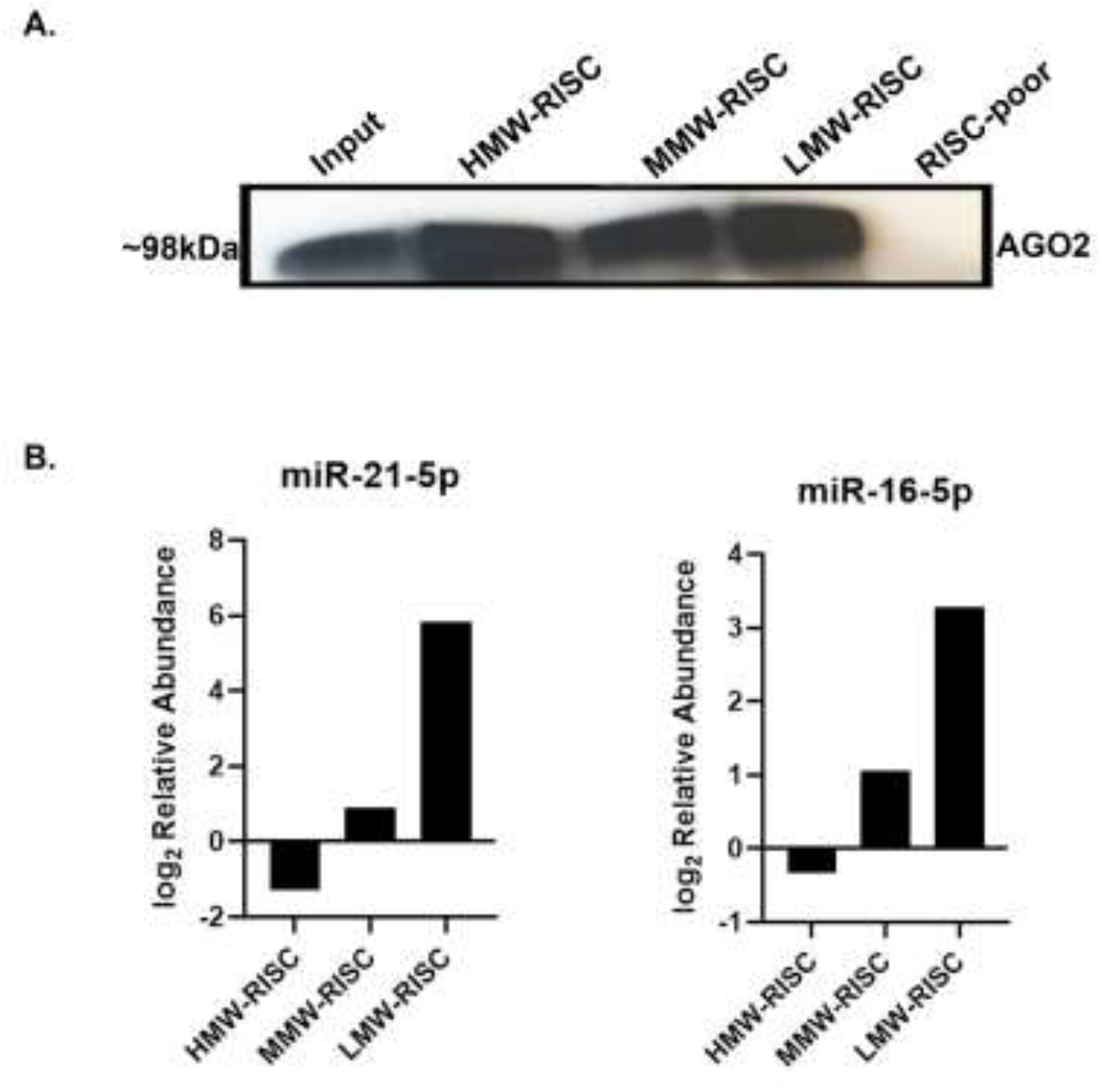
AGO2-RNA Immunoprecipitation (AGO2-RIP). **A**. AGO2 content (Western blot) from TPC-1 unfractionated input and pooled HMW-RISC, MMW-RISC, LMW-RISC, and RISC-poor SEC fractions after AGO2-RIP. **B**. miR-21-5p and miR-16-5p (qPCR) content after AGO2-RIP of TPC-1 pooled HMW-RISC, MMW-RISC, and LMW-RISC SEC fractions (normalized to RISC-poor fraction).

### Primary PBMCs Show Greater miR-21-5p Enrichment in LMW-RISC Than ATC or PTC Cell lines

Because miR-21 is one of the most extensively studied miRNAs and is reportedly overexpressed and oncogenic in various solid tumors, including thyroid cancer, the enrichment of this miRNA in inactive, LMW-RISC fractions was surprising [35]. Therefore, we assessed if the accumulation of this miRNA in mRNA-depleted fractions was ATC/PTC specific or also occurred in other cell types.

Analysis of mouse primary resting T-cell data from La Rocca et al. 2015, showed that miR-21a, which is comparable to miR-21 in humans, was present mostly in LMW-RISC fractions [28]. We observed a similar distribution of miR-21 in freshly obtained human primary PBMCs. Interestingly, however, when compared with the elution profile of miR-21-5p in SW1736, miR-21-5p LMW-RISC enrichment in human PBMCs is greater by 10-fold (Figure 7. A-B). Therefore, the size of the miR-21-5p LMW-RISC reservoir may be dependent on the cell type and may actually be smaller in thyroid cancer cell lines than in some primary cell types.

**Figure. 7.**
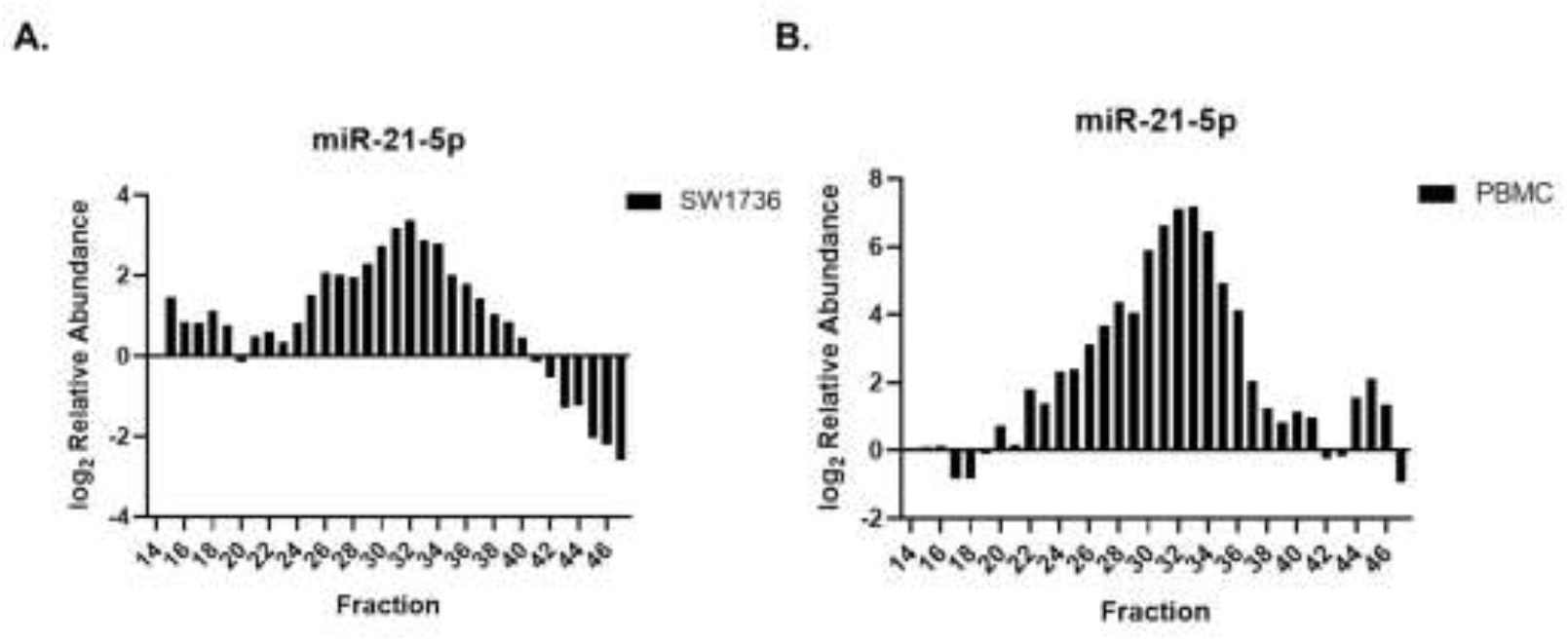
miR-21-5p across SEC fractions of SW1736 cells and peripheral blood mononuclear cells. Abundance of miR-21-5p (qPCR) across SEC fractions 14-47 relative to fraction 14 for SW1736 cells (**Panel A**) and peripheral blood mononuclear cells (PBMCs, **Panel B**).

### miR-21-5p Inhibitors Increase LMW-RISC Reservoir in SW1736

Hypothesizing that antisense inhibitors of miR-21-5p might compete with components of HMW-RISC complexes (including mRNA) and result in even greater enrichment of miR-21-5p in LMW-RISC fractions, we transfected SW1736 cells with miR-21-5p inhibitors [36]. Indeed, miR-21-5p inhibitors increased the abundance of miR-21-5p in LMW-RISC fractions and decreased HMW-RISC enrichment by greater than ten-fold (Figure 8. A-B). This result supports the conclusion that the large reservoir of miR-21-5p in LMW fractions is inactive and can be increased further by outcompeting the presumed HMW-RISC targets of miR-21-5p.

**Figure. 8.**
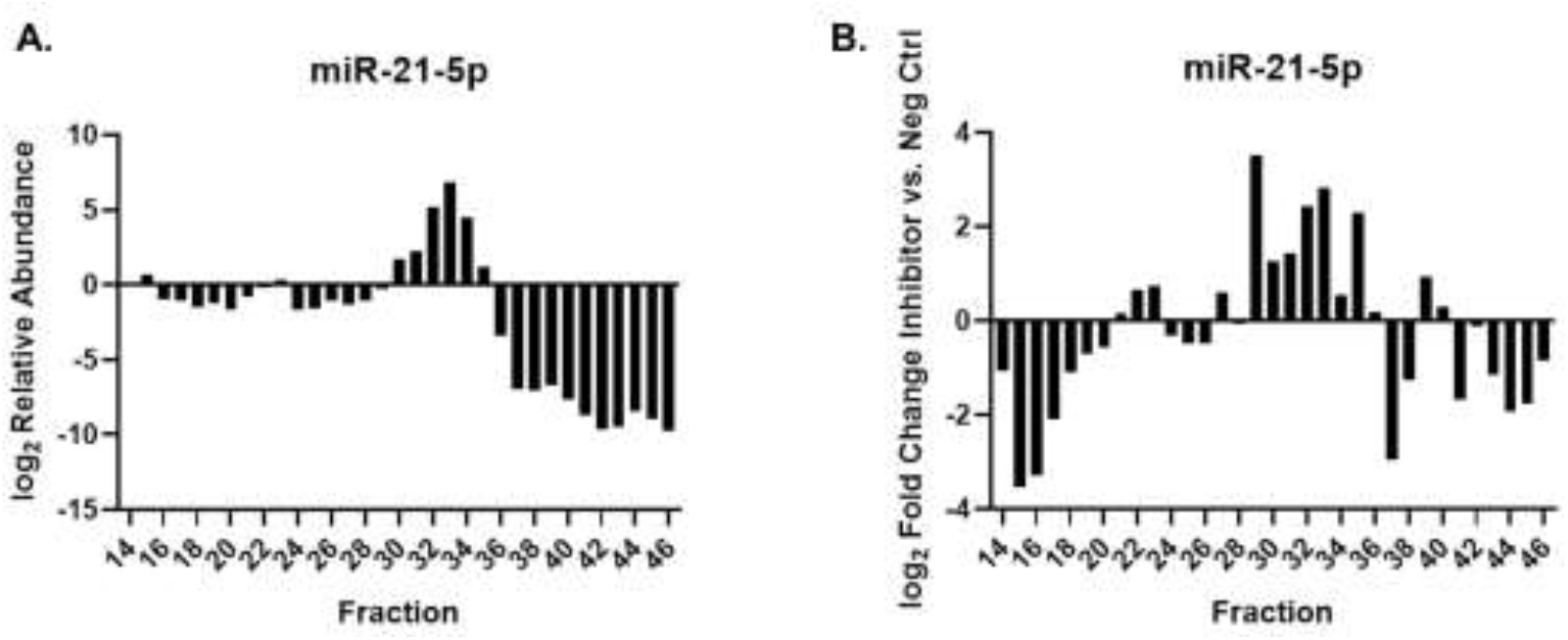
Effect of miR-21-5p antisense inhibitor on miR-21-5p distribution. SW1736 cells were transfected with a miR-21-5p antisense inhibitor. **A**. miR-21-5p abundance across SEC fractions 14-46 from transfected cells. **B**. Differential abundance of miR-21-5p in SEC fractions from miR-21-5p inhibitor-transfected cells compared with cells that received a negative control inhibitor.

### Decreased Expression of miR-21-5p in Hypoxia Lowers LMW-RISC Enrichment

In previous experiments, we observed a >2-fold decrease of miR-21-5p in SW1736 cells cultured at 2% oxygen (hypoxia) rather than in near-atmospheric levels of oxygen (unpublished results). Therefore, we investigated how lowering miR-21-5p expression levels by growth in hypoxic conditions might impact its distribution across HMW-RISC and LMW-RISC fractions. A similar distribution of miR-21-5p, which was largely enriched in LMW-RISC fractions, was observed in normoxic and hypoxic SW1736 lysates. However, the magnitude of enrichment was noticeably lower (by ~2-fold) in hypoxic SW1736. In contrast, there was only a slight decrease in HMW-RISC enrichment (<2-fold).

**Figure. 9.**
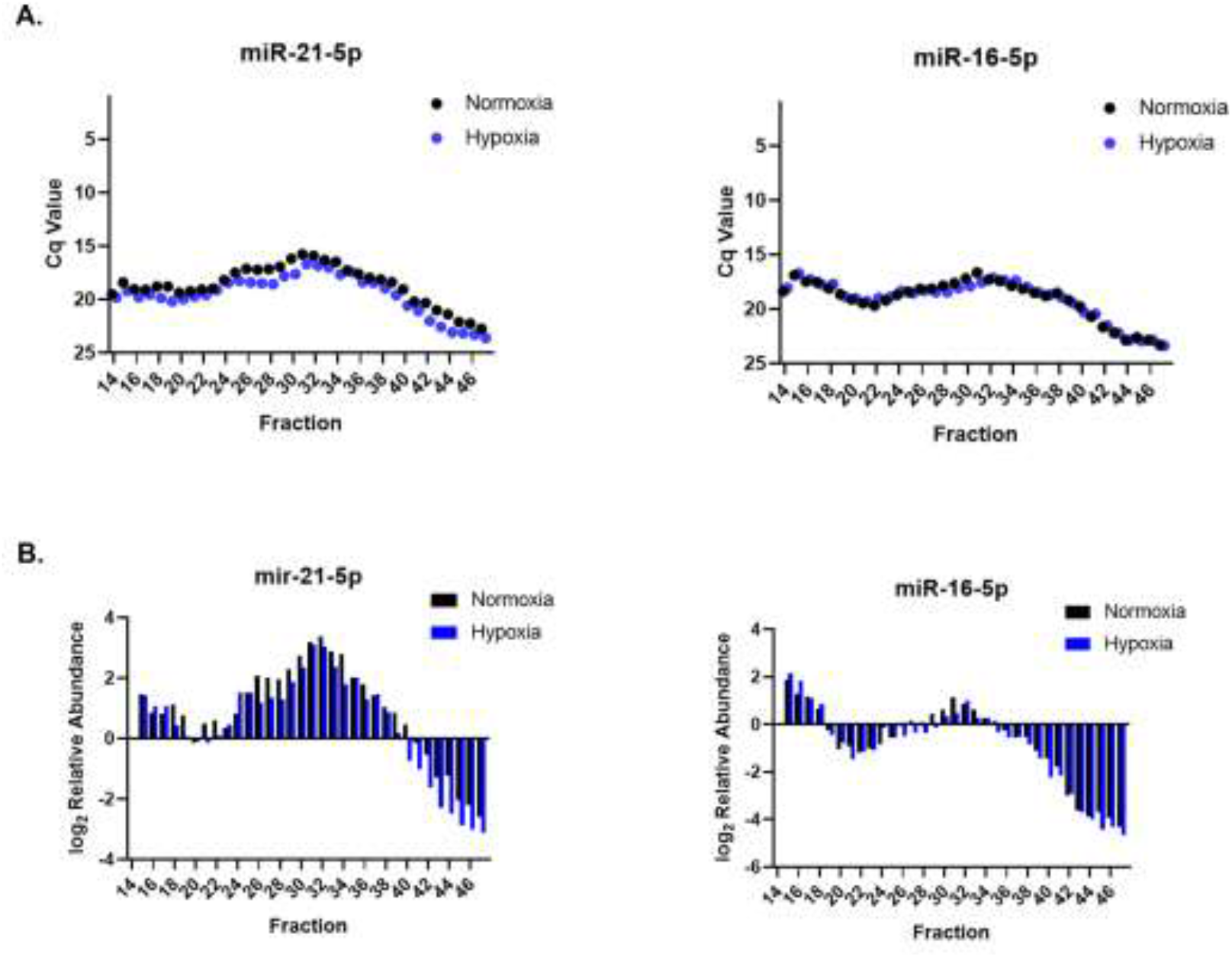
Effect of hypoxia on miR-21-5p distribution. miR-21-5p distribution across SEC fractions 14-47 after culture of SW1736 cells in hypoxic (2% O_2_) and normoxic (~21% O_2_) culture conditions. **A**. Cq values of miR-21-5p and miR-16-5p across SEC fractions 14-47 in hypoxia (blue) and normoxia (black). **B**. Relative abundance of miR-21-5p and mir-16-5p in fractions 14-47, normalized to fraction 14 (hypoxia and normoxia).

### miR-21-5p HMW-RISC Enrichment Increases with Mimic Transfection

Furthermore, to see if large quantities of miR-21 could shift enrichment from LMW-RISC to HMW-RISC fractions, SW1736 cells were transfected with single-stranded or duplex miR-21-5p mimics. With the single-stranded mimics, an overall increase in miR-21-5p abundance was observed across all SEC fractions relative to the negative control. Interestingly, miR-21 remained substantially enriched in LMW-RISC, but was also now enriched in HMW-RISC fractions by >300-fold. In contrast, miR-21-5p duplex mimics were predominantly enriched in LMW-RISC fractions.

**Figure. 10.**
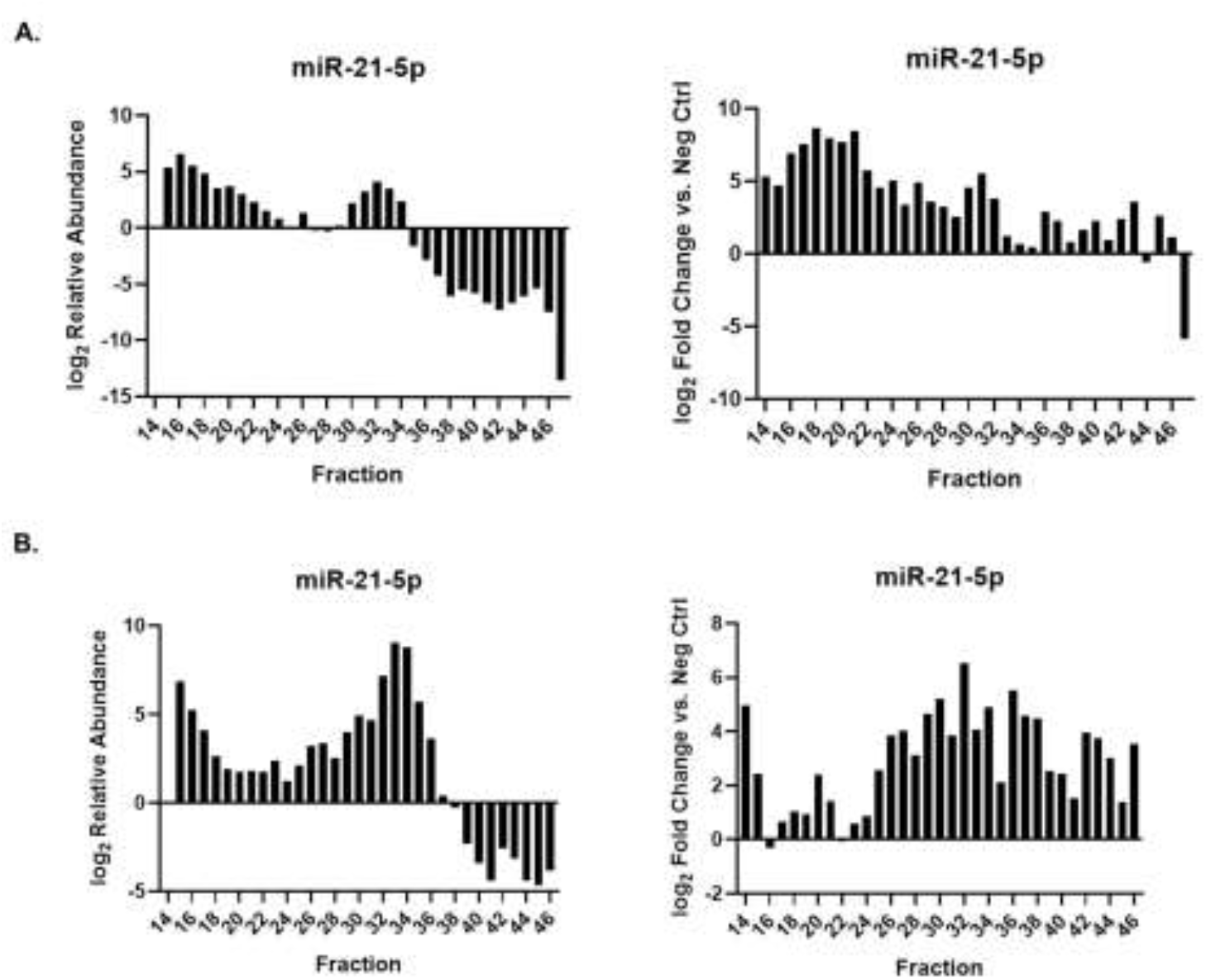
Distribution of single-stranded and duplex miR-21-5p mimics. SW1736 cells were transfected with single- or double-stranded miR-21-5p mimics or corresponding control RNAs. A. miR-21-5p detection after single-stranded mimic transfection across SEC fractions 14-47 (left panel) and relative abundance in mimic-versus control-transfected cells (right panel). B. miR-21-5p detection after double-stranded (duplex) mimic transfection across SEC fractions 14-46 (left panel) and relative abundance in mimic-versus control-transfected cells (right panel).

## DISCUSSION

Recently, it has become evident that miRNA expression differences may not be perfectly correlated with canonical mRNA repression, in part due to the influence of intercellular signaling pathways on incorporation of miRNAs into active or inactive complexes [37][38]. In this study, our results support the preferential enrichment of specific miRNAs into active, targeting RISCs or inactive small complexes. miR-21-5p is the most striking example. It is moderately abundant in the thyroid cell lines we studied, is minimally associated with target RISCs relative to other miRNAs, and is the most highly abundant miRNA in low molecular weight fractions that are not associated with mRNA.

MiR-21 is one of the most studied miRNAs [35][39][40], with numerous validated targets. It is frequently deregulated in tumors, including thyroid, and is often described as an oncogenic miRNA or oncomiR [41][42]. Numerous studies have demonstrated that aberrantly high levels of miR-21 promote tumor growth by negatively regulating the expression of tumor suppressor genes such as PTEN [43][44]. Additionally, tumors may become reliant on miR-21 in a phenomenon known as oncogene addiction, and down-regulation of miR-21 often results in tumor regression[45][46]. Effects of miR-21 have been demonstrated in both overexpression/knockout mouse models and *in vitro* transcription profiling studies[47] [46]. Because of the extensive evidence for regulatory roles of miR-21 in cancers and elsewhere, the high level of miR-21 within mRNA-poor fractions was unexpected. Indeed, our results would tend to suggest that most miR-21 copies are inactive in the cell types we examined. Because HMW-RISCs are associated with polysomes actively translating mRNA [48][49][50], it could be that the large LMW-RISC miR-21 reservoir is due to a paucity of cognate miR-21 target transcripts within HMW-RISC fractions relative to those of other miRNAs.

We also predicted that miR-21-5p might utilize a mechanism of action distinct from other miRNAs: one that does not require direct binding to a target mRNA. However, because miR-21-5p inhibitors increased the magnitude of miR-21-5p within LMW-RISC fractions and lowered levels in target complex fractions, it is evident that miR-21-5p does interact with mRNA to some degree but mostly accumulates within LMW-RISCs. The large accumulation of LMW miR-21-5p might be explained by its known low target binding affinity. Relative to other miRNAs such as let-7a, miR-21 tenuously binds to targets due to the low GC% of its seed sequence [51][52]. Hence, miR-21 may transiently interact with targets to mostly degrade mRNA through the recruitment of deadenylases or other factors, rather than remaining bound to a target and blocking the ribosome before returning to a low molecular weight state [53][54]. This might explain the pronounced biological effects on tumor growth when miR-21 expression is modulated and why it mostly exists in LMW-RISC fractions.

Although mir-21-5p is relatively abundant in both cell lines examined here, we were curious to see how exogenous miR-21-5p would be distributed between HMW-RISC and LMW-RISCs in cells using single-stranded or duplex mimics. Interestingly, increased miR-21-5p enrichment was observed in both HMW-RISC and LMW-RISC fractions with each type of mimic; yet, the single-stranded miR-21 mimics largely accumulated in HMW-RISC fractions, and duplex mimics mostly accumulated in LMW-RISC fractions. We reasoned that the single stranded mimics may not efficiently load into Argonaute proteins and that both types of mimics are likely non-specifically bound to targets, as it is known that low binding affinity of miR-21 can be supplemented by the combination of 3’end supplemental base pairing and higher expression levels [51].

Taken together, our results suggest that miR-21 may be uniquely distributed in both cancer and normal cells, with molecular and biochemical features distinct from other miRNAs. Because miR-21-5p is a promising therapeutic target for cancer, further functional characterization of this miRNA in its molecular context is crucial. The methodology described here is a useful way to assess the relative magnitude by which miR-21 is associated with HMW and LMW-RISCs and should be combined with the analysis of post pharmacological inactivation of miR-21 functional effects. Since our observations have been restricted to thyroid cell lines and PBMCs, we plan to extend our studies to normal, benign and malignant thyroid tissues in the future.

---

**Supplementary Figure. 1.**
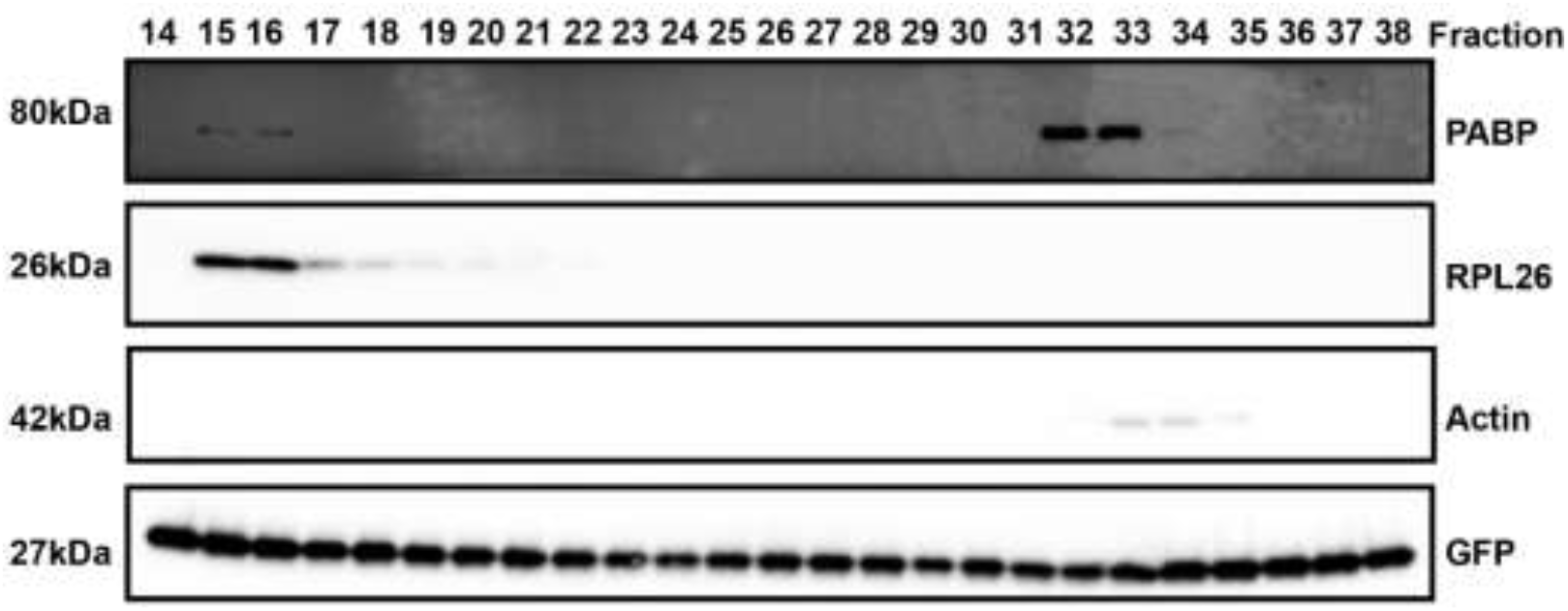
Thyroid cell line SEC elution profile. Western blot of SEC fractions 14-38. RPL26 and PABP enrichment in each fraction was analyzed to map HMW-RISCs (>2MDa). Actin complexes were also analyzed as a non-RISC associated control to confirm fractionation of lysate by SEC. GFP was spiked-in to each fraction post SEC and prior concentrating down for gel loading as a recovery control.

**Supplementary Figure. 2.**
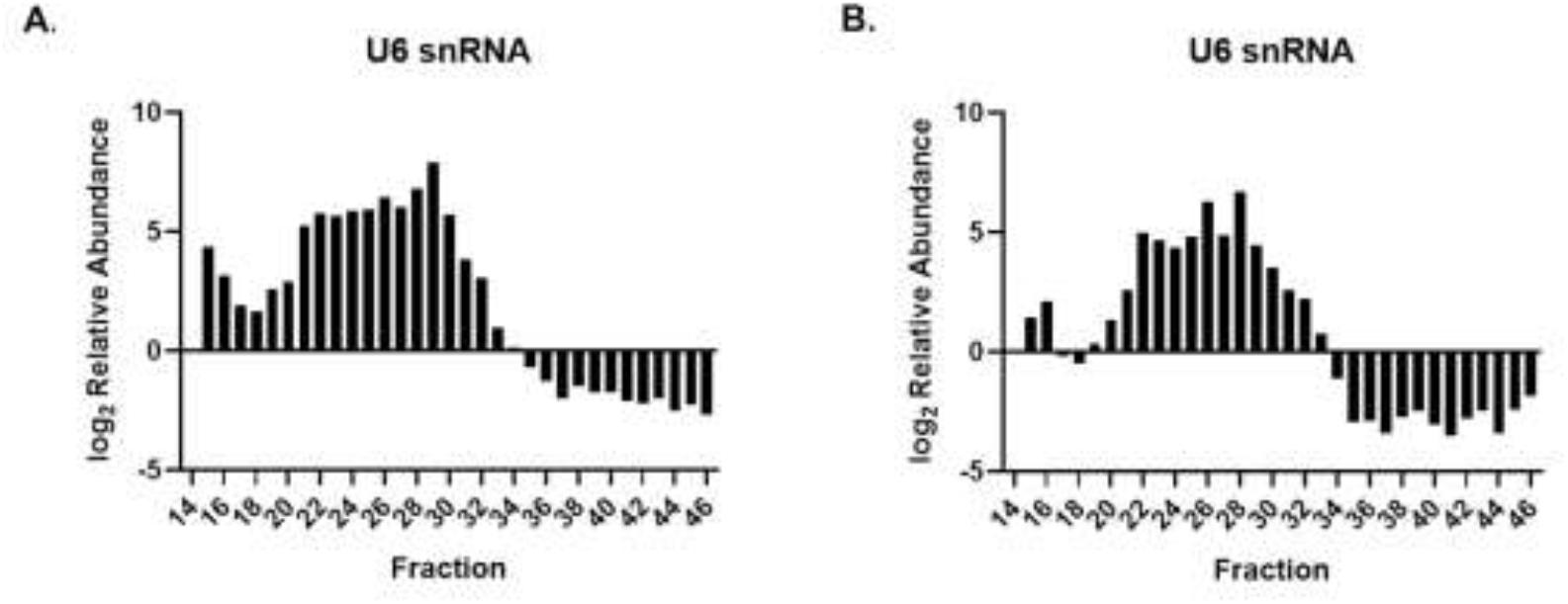
Elution profile of U6 small nuclear RNA (U6 snRNA) in thyroid cancer cell lines SW1736 and TPC-1 by RT-qPCR analysis of SEC fractions 14-47.

**Supplementary Figure. 3.**
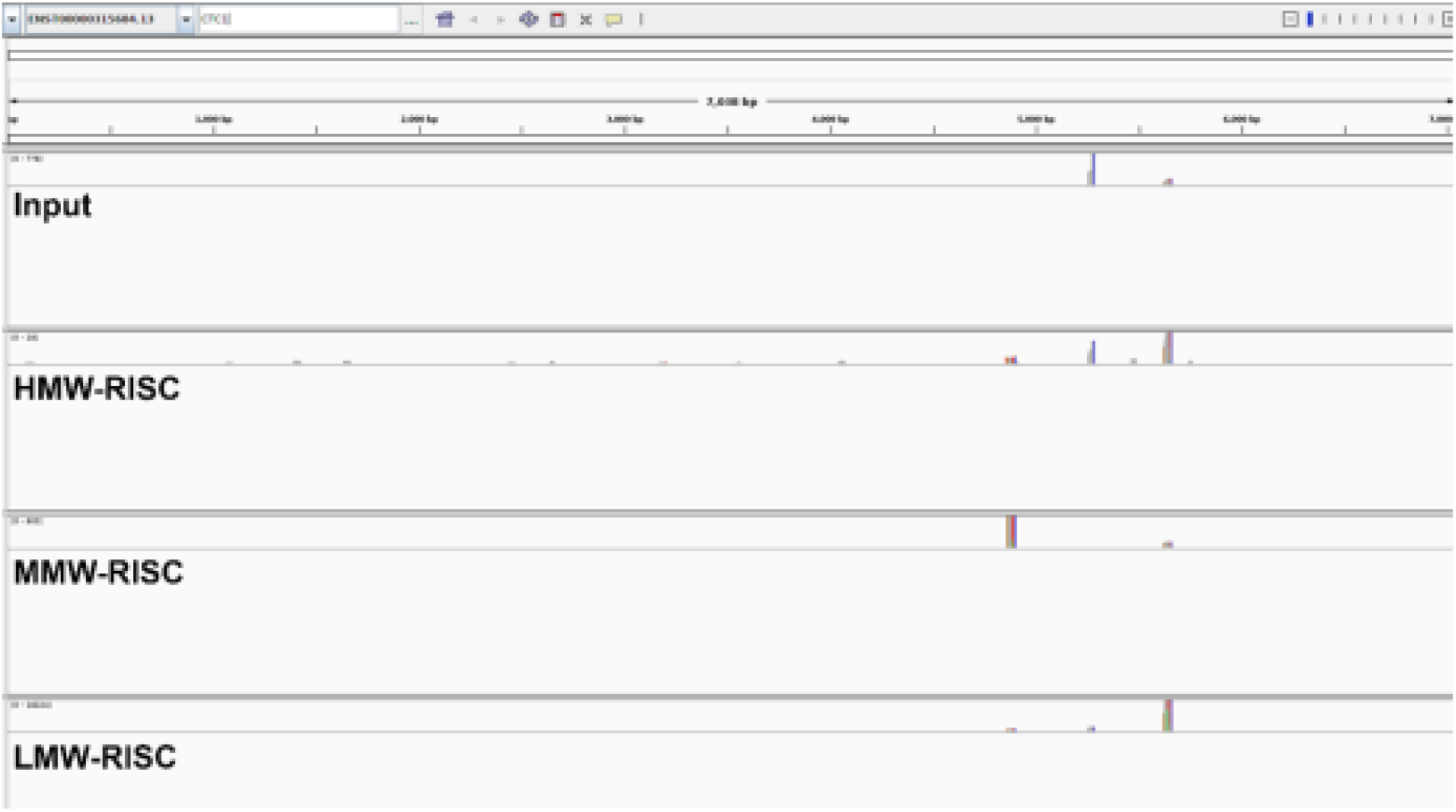
Mapping location/coverage of mRNA fragments CTC1 across pooled HMW-RISC, MMW-RISC, and LMW-RISC fractions.

**Supplementary Figure. 4.**
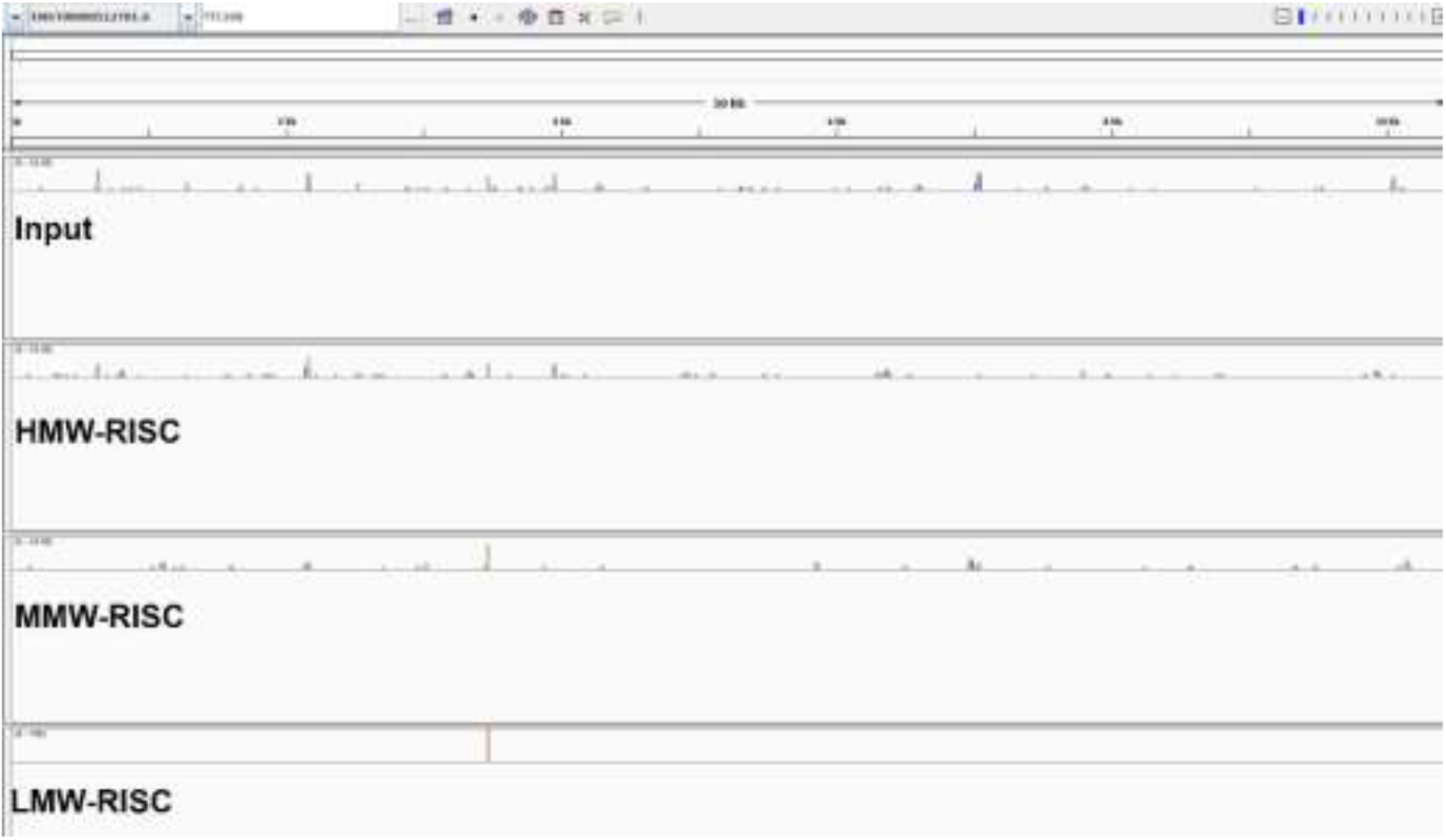
Mapping location/coverage of mRNA fragments TTC39B across pooled HMW-RISC, MMW-RISC, and LMW-RISC fractions.

**Supplementary Figure. 5.**
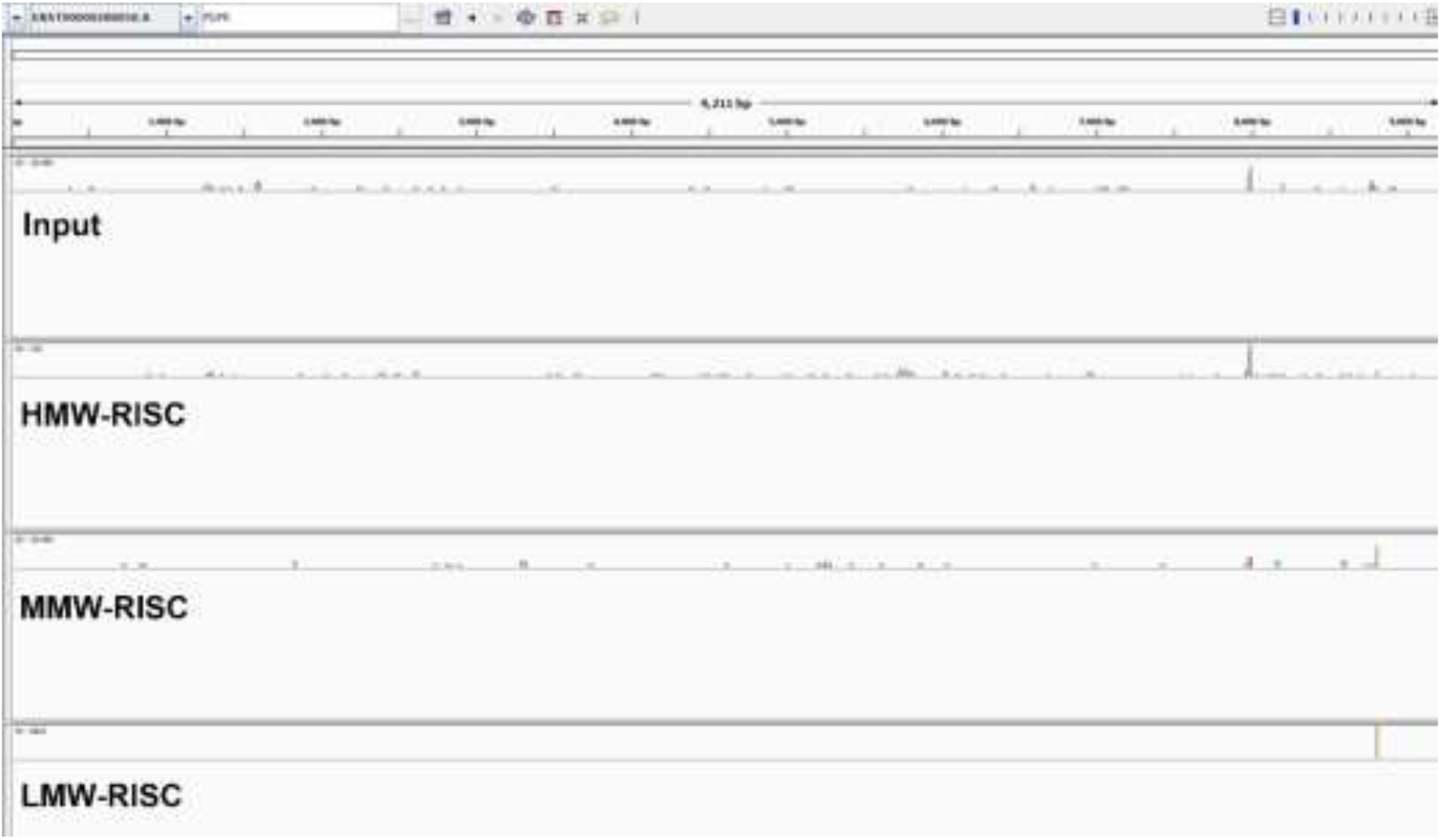
Mapping location/coverage of mRNA fragments PDPRacross pooled HMW-RISC, MMW-RISC, and LMW-RISC fractions.

**Supplementary Figure. 6.**
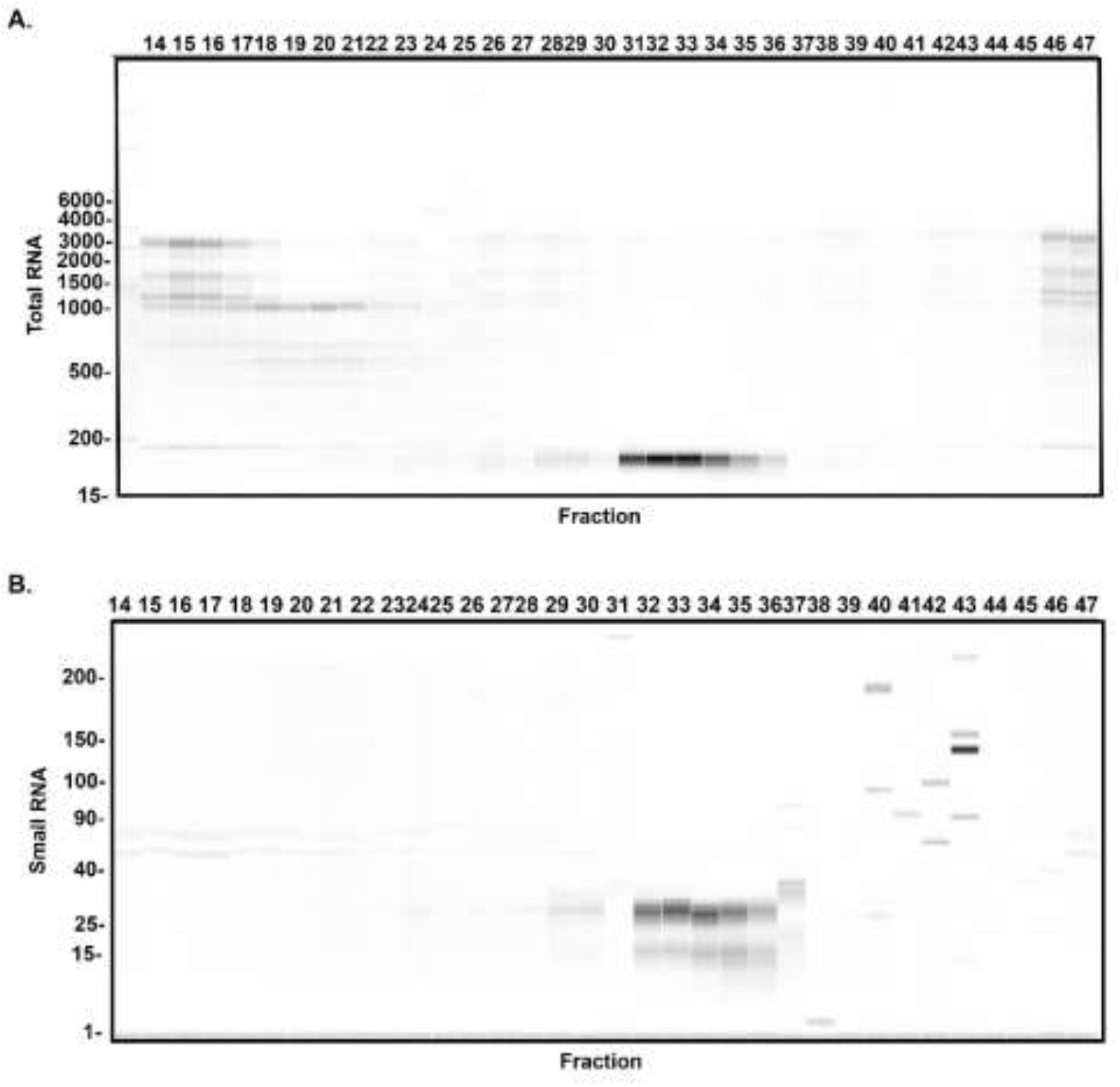
Analysis of small and total RNA across size-exclusion chromatography (SEC) fractions of thyroid cancer cell line (TPC-1). **A**. Small RNA analysis of TPC-1 SEC fractions 14-47 by fragment analyzer. **B**. Total RNA analysis of TPC-1 fractions 14-47 by fragment analyzer.

**Supplementary Figure. 7.**
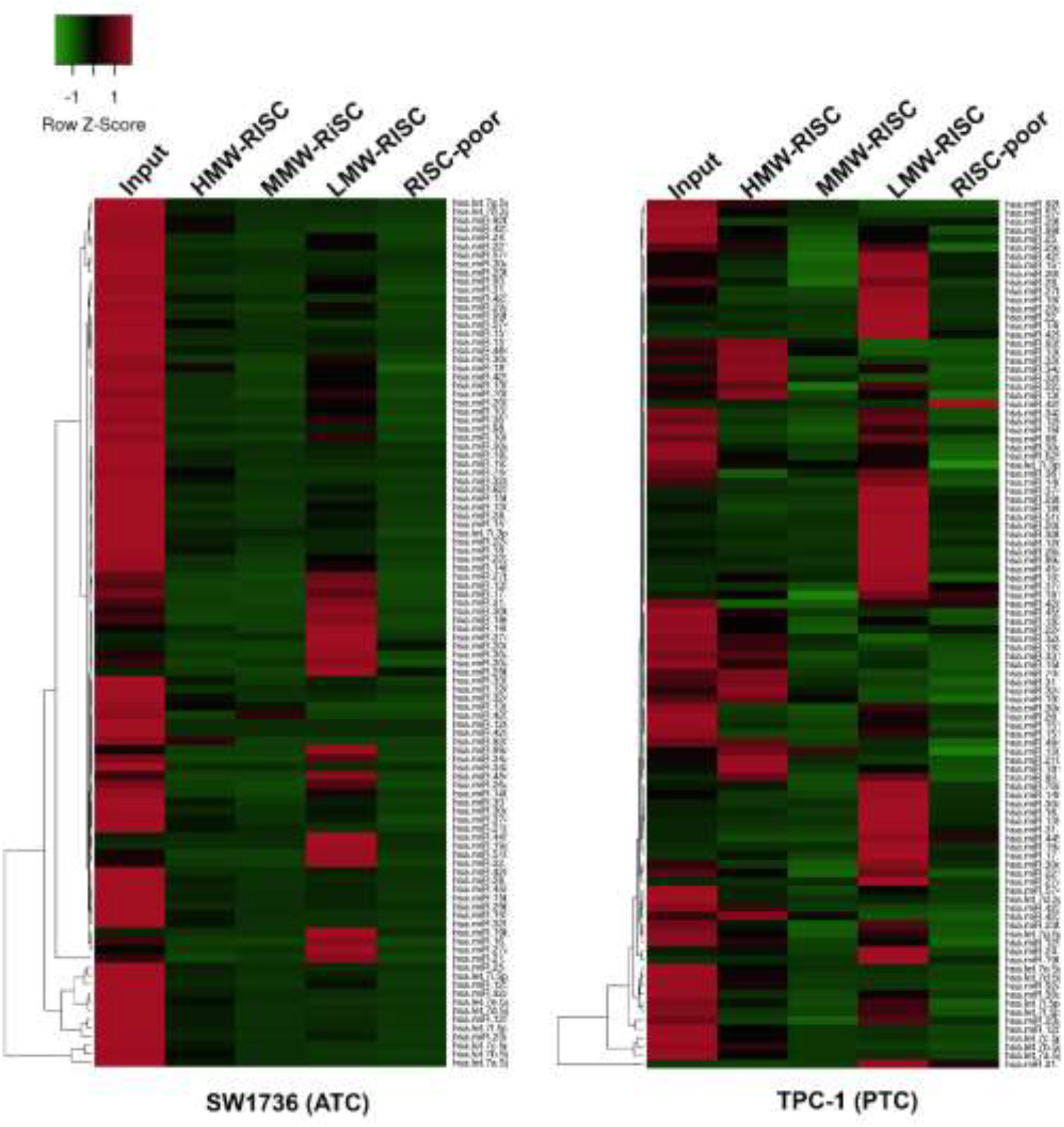
**A**. Heatmap of 101 SW1736 and TPC-1 miRNAs across HMW-RISC, MMW-RISC, and LMW-RISC fractions based on Next Generation Sequencing (NGS) log2[counts-per-million+1] (lcpm+1) for each fraction.

